# TcCARP3 modulates compartmentalized cAMP signals involved in osmoregulation, infection of mammalian cells, and colonization of the triatomine vector in the human pathogen *Trypanosoma cruzi*

**DOI:** 10.1101/2024.11.20.624561

**Authors:** Joshua Carlson, Milad Ahmed, Riley Hunter, Syeda Farjana Hoque, Joshua B. Benoit, Miguel A. Chiurillo, Noelia Lander

## Abstract

*Trypanosoma cruzi* is the causative agent of Chagas disease, a zoonotic infectious disease considered a leading cause of cardiomyopathy, disability, and premature death in the Americas. This parasite spends its life between a mammalian host and an arthropod vector, undergoing essential transitions among different developmental forms. How *T. cruzi* senses microenvironmental changes that trigger cellular responses necessary for parasite survival has remained largely unknown. Cyclic AMP (cAMP) is a universal second messenger that has been shown to regulate key cellular processes in trypanosomes, in which cyclic AMP response proteins (CARPs) have been proposed to be modulators or effectors of a PKA-independent signaling pathway. In this study we aimed to investigate the role of TcCARP3 in cAMP signaling throughout *T. cruzi* life cycle. Our results show that TcCARP3 shares a dual localization (flagellar tip and contractile vacuole complex) with adenylate cyclase 1 (TcAC1) in the main developmental stages of the parasite. We also found that TcCARP3 directly interacts with several TcACs, modulating the intracellular content of cAMP. Through generation of *TcCARP3* knockout, addback, and overexpression cell lines we showed that modulation of gene expression affects the parasite’s ability to differentiate, respond to osmotic stress, invade mammalian cells and replicate within them, and colonize the hindgut of the triatomine vector. In addition, we identified several signaling proteins interacting with TcCARP3 in what we propose are cAMP signaling microdomains. Our results unveil a key role for TcCARP3 as modulator of cAMP signals necessary for parasite differentiation and survival throughout *T. cruzi* life cycle.

**IMPORTANCE:** Cyclic AMP signaling pathways are poorly understood in the stercorarian parasite *Trypanosoma cruzi*. Specifically, the mechanisms driving the activation of TcACs in response to microenvironmental stress are completely unknown. This study unveils the role of TcCARP3 in modulating the content of cAMP through the interaction with several TcACs and putative cAMP effectors in *T. cruzi*. Particularly, TcCARP3 interacts with TcAC1 in the main developmental stages of this parasite’s life cycle, where both proteins display a dual localization pattern. These results provide new evidence supporting the compartmentalization of cAMP signals in trypanosomes. Moreover, our data unequivocally demonstrates that TcCARP3 is required for key cellular processes for parasite survival, such as response to osmotic stress, host cell invasion, intracellular replication, and the ability to colonize the hindgut of the triatomine vector. In summary, we found that TcCARP3 is an adenylate cyclase regulator, necessary for the life cycle progression of *T. cruzi*.

## INTRODUCTION

*Trypanosoma cruzi* is the protozoan parasite that causes Chagas disease, one of the neglected tropical diseases of greatest public health importance in the Americas. According to the most recent data available from the World Health Organization (WHO), an estimate of 7 million people are affected by this debilitating disease, with 70 million people at risk of infection (1). Many people with Chagas disease are not even aware of their condition, as initially the disease show non-specific flu-like symptoms and patients resume normal daily life, but the infection persists. It is not until decades later that roughly a third of patients will develop serious health conditions such as cardiac, gastrointestinal, or neurological complications that can result in death (2, 3). Currently, only two drugs are available to treat Chagas disease while patients are in the acute phase of infection, but once it progresses to the chronic phase, these drugs become ineffective (4). While endemic to 21 Latin American countries, an increasing number of Chagas disease cases has been reported in many non-endemic regions including the United States, Canada, Europe, the Middle East, Asia, and Australia (3). The main driving forces behind this spread in recent decades are global human migrations from endemic to non-endemic countries, and vector colonization of non-rural areas as a consequence of climate change (1, 4). As Chagas disease becomes a global health problem, the development of alternative and more efficient strategies to diagnose and treat *T. cruzi* infections is urgently needed. Understanding this parasite’s biology is crucial for the rational development of new antiparasitic interventions.

*T. cruzi* is a stercorarian parasite with a digenetic life cycle alternating between an arthropod vector (a triatomine bug) and a mammalian host. The natural transmission of *T. cruzi* to people is from the feces of an infected triatomine, commonly known as kissing bugs. Infective metacyclic trypomastigotes (MTs), are present in the urine and feces of the vector and are transmitted to the mammalian host via skin wound or mucous membranes. Once inside the organism, MTs can invade any nucleated cell and differentiate into the intracellular amastigote. After several rounds of replication, amastigotes differentiate into infective cell-derived trypomastigotes that are eventually released into the bloodstream. These trypomastigotes can then invade other host cells or can be taken up by another triatomine bug. Once inside the vector, trypomastigotes differentiate into the proliferative epimastigotes in the midgut. Over the course of several weeks epimastigotes migrate to the hindgut of the triatomine bug, where they differentiate into infective MTs, in a process known as metacyclogenesis [reviewed by (5)]. Throughout its life cycle, *T. cruzi* encounters significant microenvironmental changes, including drastic fluctuations in temperature, pH, nutrient availability and composition, and osmolarity [Reviewed by (6, 7)]. The underlying mechanisms of how *T. cruzi* senses these environmental changes and triggers specific cellular responses that drive developmental transitions are still poorly understood.

As in many eukaryotic organisms, *T. cruzi* relies on cAMP signaling to mediate cellular responses to external stimuli (7, 8). However, to date this pathway has been better characterized in the mammalian system, where the nature and function of specific proteins determining the spatiotemporal dynamics of the signals has been well described (9, 10). Canonically, an external stimulus is received by a G protein coupled receptor (GPCR) and that signal is transduced to adenylate cyclases (ACs), enzymes responsible for the catalytic conversion of ATP into cAMP. This second messenger interacts with effector proteins such as protein kinase A (PKA), exchange protein activated by cAMP (EPAC), or cyclic nucleotide-gated (CNG) ion channels. The signal is then abolished with the degradation of cAMP into AMP by phosphodiesterases (PDEs) (11). In trypanosomes, catalytically active ACs and PDEs have been well characterized (12-19), but the canonical effectors of cAMP are either absent or not responsive to cAMP in these parasites (20, 21). Furthermore, genes encoding GPCRs are absent in the genomes of trypanosomatids (22, 23), leading to fundamental questions such as what mechanism drives the activation of ACs in these parasites, and what downstream effectors are modulated by cAMP upon its synthesis (24-26). In *T. cruzi*, ACs comprise a multigene family that we have classified in 5 groups of putative receptor-type adenylate cyclases (AC I-V) (16). In this parasite, cAMP signaling has been specifically linked to the cellular processes of cell adhesion (16), metacyclogenesis (16, 27-31), and response to osmotic stress (15, 16, 32-35), but further research is needed to understand the molecular mechanisms driving these responses. A promising group of proteins that includes putative cAMP effectors in trypanosomatids are known as cyclic AMP response proteins (CARPs). CARPs are kinetoplastid-specific proteins that were first identified in *T. brucei* through a genome-wide RNAi screening for resistance to lethal concentrations of PDE inhibitors (36). One of these proteins, CARP3, has been characterized in *T. brucei* (TbCARP3) as a multi-adenylate cyclase modulator involved in Social Motility (SoMo) and colonization of the insect vector (37, 38). We recently found that the *T. cruzi* homolog (TcCARP3) shows a peculiar dual localization pattern in the flagellar distal domain (flagellar tip) and the contractile vacuole complex (CVC) of the parasite (16). These two compartments are directly involved in cell adhesion/metacyclogenesis and response to osmotic stress, respectively (5, 32, 34, 35, 39-41). In addition, these processes have previously been linked to cAMP signaling (15, 27, 29-31, 33, 34, 42). Furthermore, TcCARP3 localization mirrored that of the catalytically active TcAC1, and their interaction was demonstrated through immunoprecipitation and mass spectrometry analysis using TcAC1 as bait (16). Considering the localization of TcCARP3 and TcAC1 in these two subcellular compartments, their peculiar presence in the CVC, and the stark differences in the biology of *T. brucei* (salivaria) and *T. cruzi* (stercoraria) parasites, we aimed to investigate the role of TcCARP3 in cAMP signaling throughout *T. cruzi* life cycle. In this study we modulated the expression of TcCARP3 through generation of mutant cell lines and evaluated their phenotype in different developmental stages, *in vitro* and *in vivo*. Our results shed light on the role of TcCARP3 in cAMP signaling and its protein interactors in two distinct subcellular compartments, leading to cellular responses necessary for parasite survival and transmission during the progression of *T. cruzi* life cycle.

## RESULTS

### CARP3 is a *Trypanosoma*-specific protein with dual localization in *T. cruzi*

*TcCARP3* (TriTrypDB gene ID: TcYC6_0045920) is a 1548-bp single copy gene annotated as hypothetical protein on chromosome 16 of *T. cruzi* Y C6 genome (43, 44). The predicted protein has a molecular weight of 58.16 kDa and is 515 amino acids in length, with a high confidence predicted post-translational modification of myristoylation occurring on amino acid number 2, glycine, immediately following the start methionine (45). TcCARP3 tertiary structure also has a predicted TPR like tetratricopeptide-like helical domain spanning from amino acids 13-155, as predicted by Interpro (IPR011990) in TriTrypDB (44, 46) (Fig. 1A). This domain has important implications in mediating protein-protein interactions and the assembly of multi-protein complexes in a wide range of proteins from a diverse set of organisms (47-50). *TcCARP3* shares 62.47% nucleotide sequence identity with its ortholog in *T. brucei, TbCARP3* (TriTrypDB gene ID: Tb427.07.5340) and 50.09% identity at the amino acid level, with 68.41% protein similarity. Conversely, *CARP3* orthologs are absent in *Leishmania spp*. We have previously shown that TcCARP3 and TcAC1 (TriTrypDB gene ID: TcYC6_0015740) co-localize in 2 different compartments of *T. cruzi* epimastigotes: the flagellar tip and the CVC (16). To further confirm TcCARP3’s dual localization in this stage we endogenously tagged TcCARP3 with a C-terminal 3xTy1 tag as described in *Materials and methods*. The expression and dual localization of TcCARP3 was confirm using this cell line (Fig. S1). Then, using a dually tagged cell line expressing TcCARP3-3xc-Myc and TcAC1-3xHA (16), we analyzed the localization of both proteins by immunofluorescence analysis (IFA) in the four main developmental stages of *T. cruzi*. IFAs were done under hypoosmotic conditions to better visualize the central vacuole of the CVC. Our results indicate that these proteins co-localize in all developmental stages (epimastigotes, metacyclic trypomastigotes, amastigotes and cell-derived trypomastigotes), showing the previously described dual localization pattern in all of them, except in metacyclic trypomastigotes, where both proteins localized to the tip of the flagellum only (Fig. 1B). The flagellar tip localization of TcCARP3 has been previously reported in the intracellular amastigote stage (51), but we also observed it in the CVC of *T. cruzi* epimastigotes (16). Here we have confirmed this dual localization pattern in the mammalian stages of the parasite.

**Figure 1.**
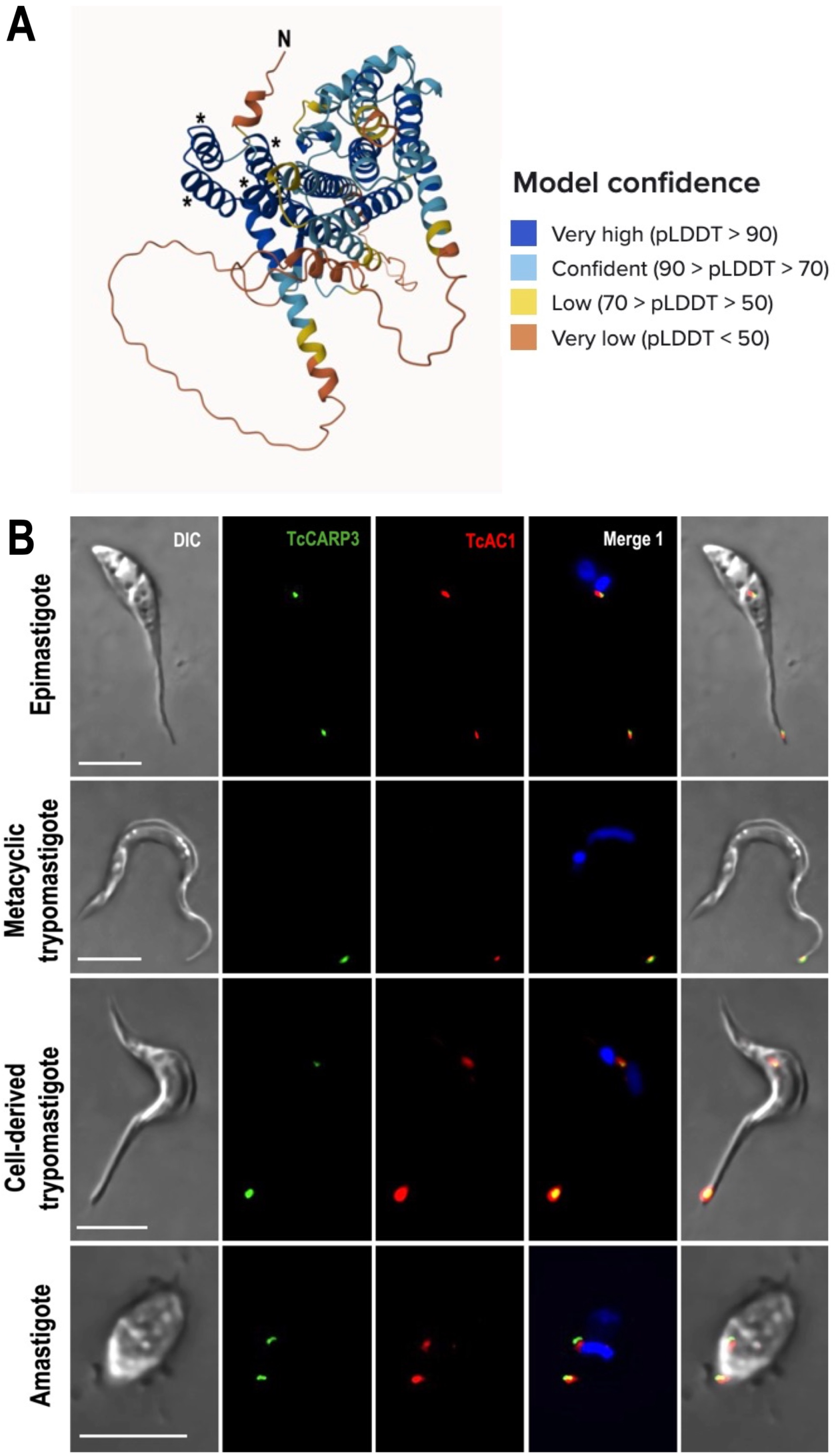
TcCARP3 predicted structure and co-localization of TcCARP3 and TcAC1 in different developmental stages of *T. cruzi.* **[A]** TcCARP3 structure predicted with AlphaFold. Model confidence using per-residue confidence score (pLDDT) is indicated by color-code. Regions below 50 pLDDT may be unstructured in isolation. Asterisks indicate the four alpha helices predicted to constitute the Tetratricopeptide-like helical domain at the N-terminus of TcCARP3. **[B]** IFAs were performed using TcAC1-3xHA/TcCARP3-3xc-Myc dually tagged cell line under hypoosmotic stress. Images from left to right show DIC, TcCARP3 (green), TcAC1 (red), TcCARP3 and TcAC1 merged (yellow) with DAPI (blue), and with DIC. DAPI was used to stain the nucleus and kinetoplast. Scale bars: 5 μm.

### TcCARP3 is involved in growth and metacyclogenesis but not in cell adhesion

To analyze the effect of *TcCARP3 gene* ablation if different developmental forms, we generated a *TcCARP3* knockout mutant by CRISPR/Cas9 (*TcCARP3*-KO), as described in *Materials and Methods* (Fig. 2A). After confirmation of this mutant genotype (Fig. 2B, C) clonal populations were obtained by serial dilutions and the resulting clones were verified by PCR (Fig. S2). The *TcCARP3* gene was then added back by cloning its ORF into pTREXh-2xTy1 expression vector and transfecting a clonal population of *TcCARP3*-KO parasites to generate the *TcCARP3* addback cell line (*TcCARP3*-AB). Expression of TcCARP3-2xTy1 in *TcCARP3*-AB parasites was verified by western blot analysis (Fig. 2D). A growth curve was performed using the parental cell line (T7/Cas9) as control, *TcCARP3*-KO, and *TcCARP3*-AB parasites (Fig. 3A). The growth rate in LIT medium was examined during the exponential phase (days 3-6), resulting in a significantly lower growth of *TcCARP3*-KO epimastigotes compared to control cells, while the normal phenotype was partially rescued in *TcCARP3*-AB parasites. A TcCARP3 overexpressing cell line (*TcCARP3*-OE) was obtained as described in *Materials and Methods*. Briefly, the ORF of *TcCARP3* was cloned into pTREXn-3xHA (16) and used to transfect wild type (WT) epimastigotes. Expression of TcCARP3-3xHA in a clonal population was confirmed by western blot analysis (Fig. S3A). We then evaluated the growth of *TcCARP3*-OE epimastigotes in LIT medium, compared to that of the pTREXn-3xHA empty vector (EV) control, and no significant difference was observed (Fig. S3B). TcCARP3 mutant cell lines were also used to evaluate metacyclogenesis, a process that is essential for parasite development within the triatomine vector and further transmission to mammalian hosts. *In vitro* metacyclogenesis was performed by incubating epimastigotes in triatomine artificial urine (TAU) to simulate the conditions in the hindgut of the triatomine bug. Then, we evaluated the percentage of metacyclic trypomastigotes by fluorescence microscopy upon DAPI staining. Interestingly, *TcCARP3*-KO parasites showed a significantly higher percentage of metacyclic trypomastigotes compared to the control, and this phenotype was rescued by *TcCARP3*-AB parasites (Fig. 3B). We also performed *in vitro* metacyclogenesis for *TcCARP3*-OE and control parasites and found no significant differences among them (Fig. S3C). In the kissing bug, metacyclogenesis is preceded by attachment of the parasite through the flagellar tip to the hindgut cuticle (5, 39). To test the ability of *TcCARP3*-KO parasites to adhere during this process we performed an *in vitro* adhesion assay by placing epimastigotes in TAU3AAG and counting the number of cells in the supernatant with a Neubauer chamber at different time points. Surprisingly, we did not observe a significant difference in the adhesion capacity of *TcCARP3*-KO, *TcCARP3*-AB, and control parasites (Fig. 3C), indicating that the metacyclogenesis phenotype observed in these mutants is independent of their adhesion phenotype.

**Figure 2.**
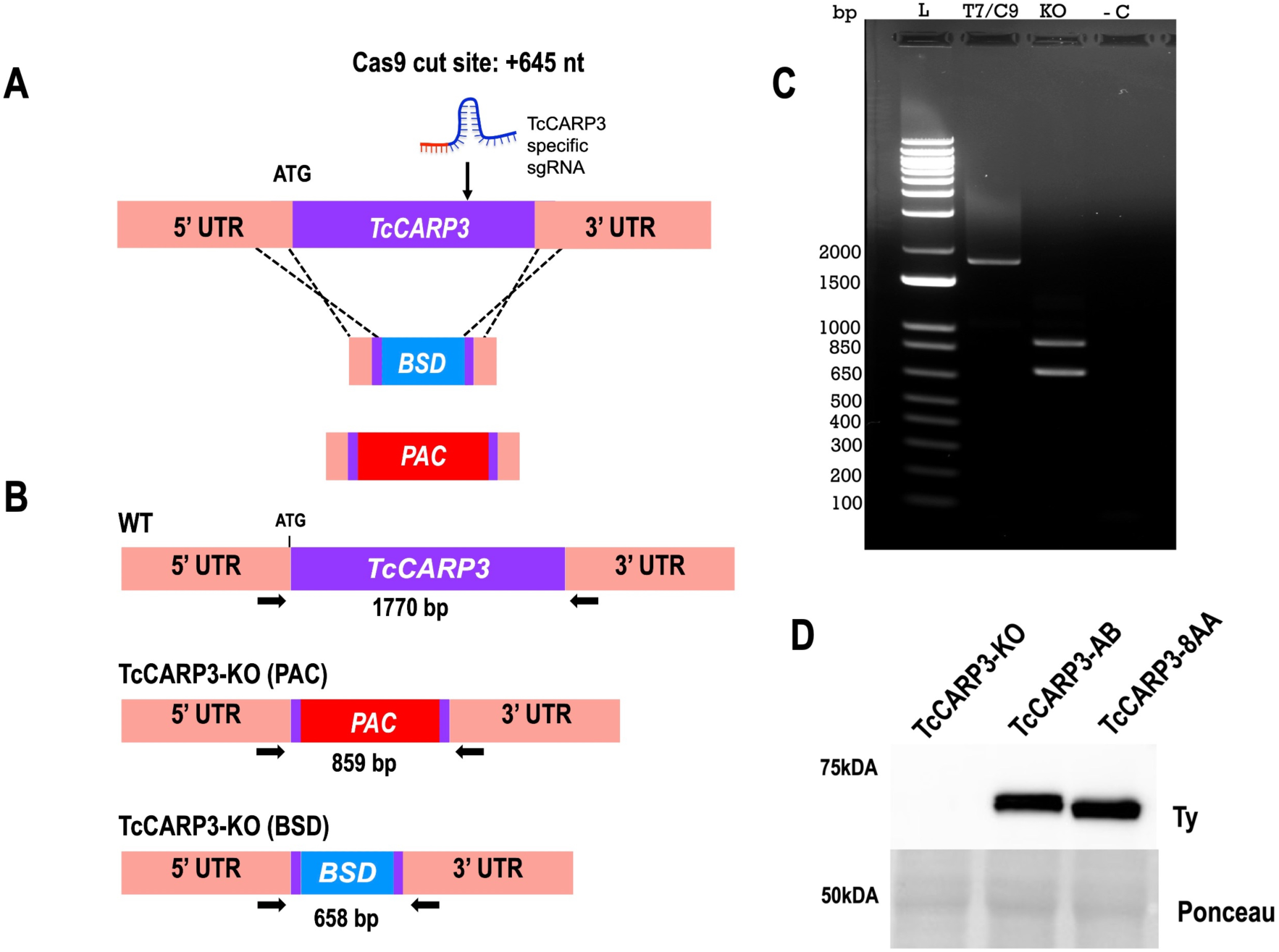
Generation of *TcCARP3* knockout and addback cell lines. **[A]** Schematic representation of CRISPR/Cas9-mediated knockout strategy for *TcCARP3*. **[B]** Predicted sizes of PCR products using parental or *TcCARP3*-KO gDNA. **[C]** After selection of clones and gDNA extraction, PCR was performed to verify the genotype, and products were resolved in a 1% agarose gel. **[D]** Western blot analysis confirming expression of TcCARP3 in the *TcCARP3*-AB (61.15kDa) and *TcCARP3*-8AA (61.05kDa) cell lines. Ponceau red staining was used as loading control.

**Figure 3.**
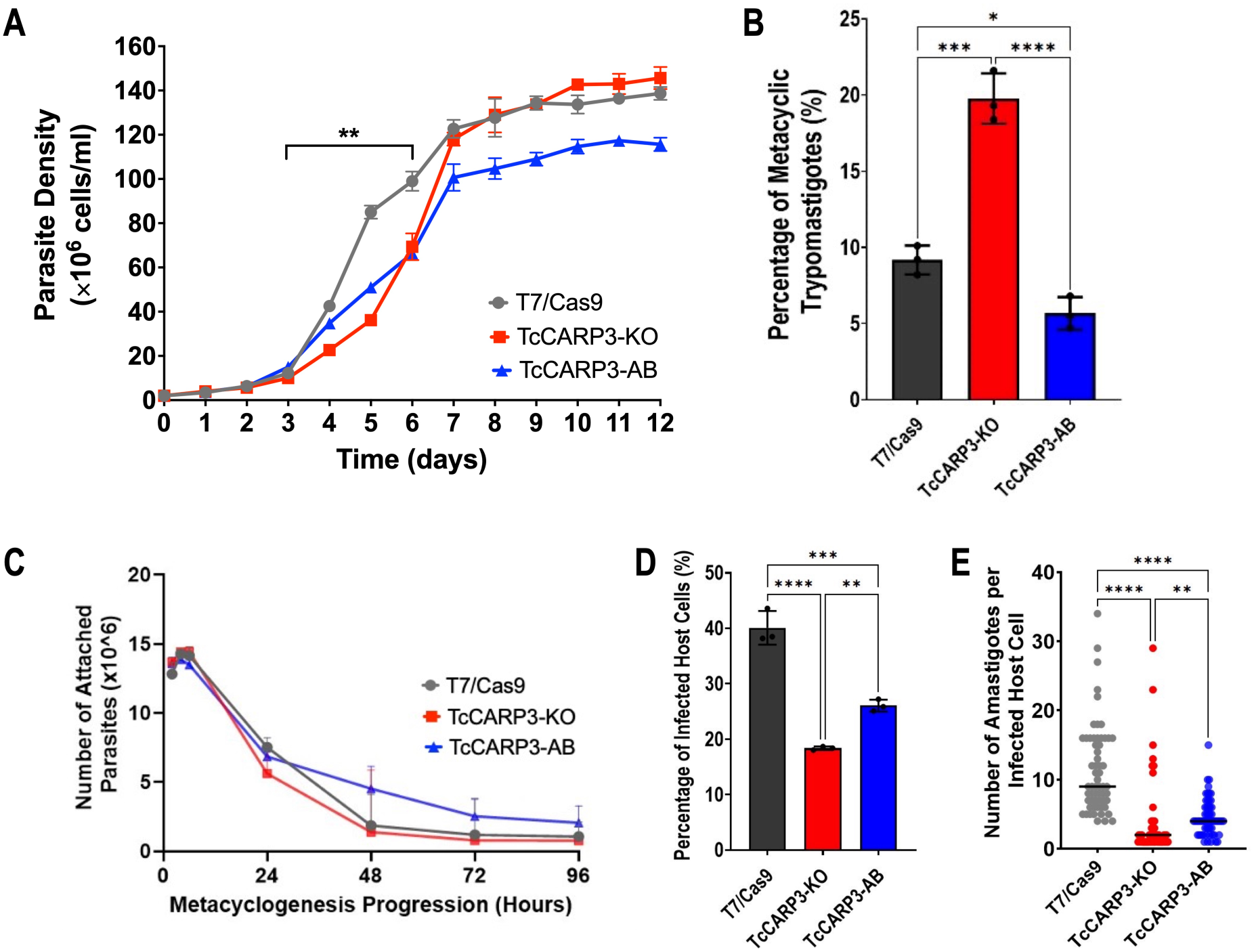
Phenotype of *TcCARP3* mutants. **[A]** Growth of T7/Cas9, *TcCARP3*-KO, and *TcCARP3*-AB epimastigotes in LIT medium. Growth rate was analyzed during exponential phase of the curve (Days 3-6). Values are means ± S.D., n=3. One-way ANOVA with Dunnett’s multiple comparisons. **[B]** Metacyclogenesis *in vitro* of T7/Cas9, *TcCARP3*-KO, and *TcCARP3*-AB parasites. Values are means ± S.D., n=3. One way ANOVA with Tukey’s multiple comparisons. **[C]** Adhesion assay with T7/Cas9, *TcCARP3*-KO, and *TcCARP3*-AB parasites. No significant differences were observed at any time point. Values are means ± S.D., n=3. Two-way ANOVA with Dunnett’s multiple comparisons. **[D]** Percentage of infected host cells 24 h post infection with T7/Cas9, *TcCARP3*-KO, and *TcCARP3*-AB trypomastigotes. Values are means ± S.D., n=3. One way ANOVA with Tukey’s multiple comparisons. **[E]** Number of intracellular amastigotes per infected host cell 72 h post infection with T7/Cas9, *TcCARP3*-KO, and *TcCARP3*-AB trypomastigotes. Black line indicates median value per cell line, n=60. Kruskal-Wallis test with Dunn’s multiple comparisons. *P< 0.05, **P< 0.01, ***P< 0.001, ****P< 0.0001.

### TcCARP3 plays a role in host cell invasion and intracellular replication

To progress in their life cycle, *T. cruzi* metacyclic trypomastigotes invade mammalian host cells and intracellularly differentiate into replicative amastigotes. We previously found that cAMP modulates the ability of parasites to invade host cells and replicate within them (16). Based on this observation, we evaluated the invasion and intracellular replication phenotype of TcCARP3 mutants using standard methods. Cell-derived trypomastigotes from T7/Cas9, *TcCARP3*-KO and *TcCARP3*-AB cell lines were used to infect human foreskin fibroblasts (hFFs). These infected cells were fixed and mounted onto slides with DAPI. The number of infected host cells (24 h post infection) and the number of amastigotes per infected cell (72 h post infection) were determined by fluorescence microscopy. The percentage of host cells infected with *TcCARP3*-KO parasites was significantly lower than those infected with control trypomastigotes, while *TcCARP3*-AB parasites partially restored the normal phenotype (Fig. 3D). In addition, the intracellular replication *TcCARP3*-KO amastigotes was also hindered, showing a significantly lower number of intracellular amastigotes per infected cell compared to the T7/Cas9 control. Again, this phenotype was partially rescued by *TcCARP3*-AB parasites (Fig. 3E). Our results indicate that TcCARP3 is required for invasion of mammalian host cells by *T. cruzi* cell-derived trypomastigotes and for the replication of amastigotes within the cell.

### Ablation of a predicted myristoylation signal does not alter TcCARP3 dual localization

Myristoylation is a post-translational modification (PTM) that involves the addition of a fourteen-carbon unsaturated fatty acid chain to a subset of N-terminal glycine residues. This PTM has important implications in membrane association and localization of proteins (52). TcCARP3 exhibits two predicted myristoylation sites, the glycine residues at the second (highest score, in a consensus sequence) and eighth position in the N-terminal end of the protein (Fig. S4A). To evaluate if this predicted myristoylation signal was required for TcCARP3 dual localization in *T. cruzi* we transfected *TcCARP3*-KO epimastigotes with a pTREXh-2xTy1 vector containing a truncated version of *TcCARP3*, called *TcCARP3*-8AA, where the two glycine residues encoded within the first eight amino acids were deleted. Expression of TcCARP3-8AA was confirmed by western blot analysis (Fig. 2D). These parasites showed the same dual localization pattern (the flagellar tip and CVC) as in the wild type addback, *TcCARP3*-AB (Fig. S4B, C), suggesting that the predicted myristoylation signal of TcCARP3 is not required for its dual localization in *T. cruzi* epimastigotes.

### Modulation of TcCARP3 expression affects the cAMP content in *TcCARP3* mutants

To test if the interaction between TcCARP3 and TcAC1 modulates the activity of the latter, we evaluated the levels of cAMP in *TcCARP3* mutant parasites using the luminescent cAMP Glo-Assay (Promega), as described in *Materials and Methods*. Our results indicate that *TcCARP3*-KO parasites exhibit a significantly lower content of cAMP than that of control cells. Interestingly, the relative levels of cAMP in *TcCARP3*-AB parasites were significantly higher than those of the T7/Cas9 control and *TcCARP3*-KO parasites (Fig. 4A). We also analyzed the effect of TcCARP3 overexpression on total cAMP content and found that *TcCARP3*-OE parasites showed significantly higher levels of cAMP compared to those of the empty vector control (Fig. 4B). These results indicate that modulation TcCARP3 expression in *T. cruzi* epimastigotes impact their relative content of cAMP, possibly due to the regulation of TcAC1 activity by TcCARP3.

**Figure 4.**
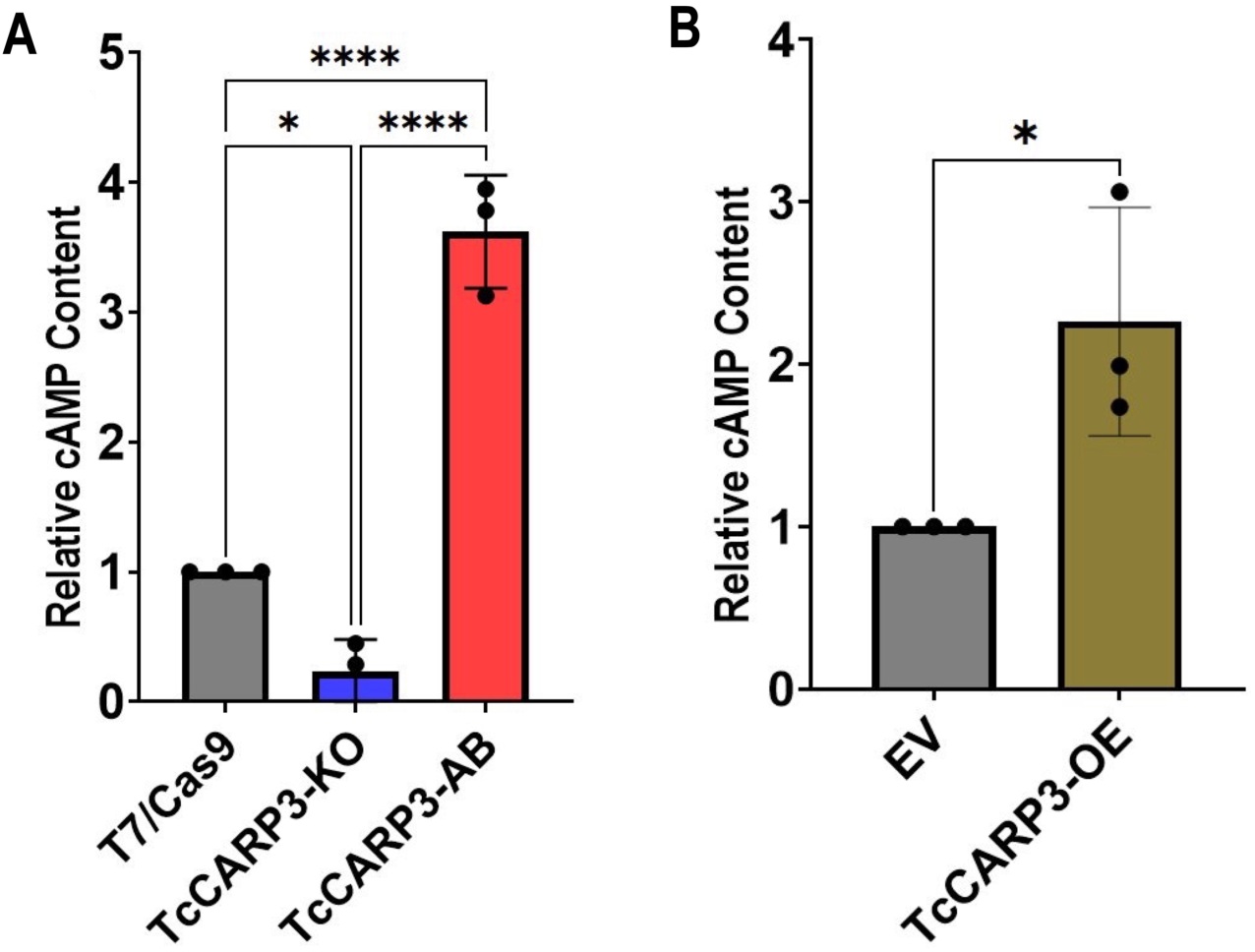
Intracellular cAMP content in *TcCARP3* mutants. **[A]** For T7/Cas9, *TcCARP3*-KO, and *TcCARP3*-AB *P< 0.05, ****P< 0.0001 (One way ANOVA with Tukey’s multiple comparisons) ± S.D. n=3. **[B]** For *TcCARP3*-OE and Empty Vector (EV) *P< 0.05 (Student’s t test) ± S.D. n=3.

### TcCARP3 plays a role in the osmoregulatory capacity of *T. cruzi* epimastigotes

The role of cAMP in the ability of *T. cruzi* epimastigotes to respond to hypoosmotic stress through a process called regulatory volume decrease (RVD) has been previously reported (21, 35-37). Since TcCARP3 co-localizes with TcAC1 in the contractile vacuole complex, an organelle specialized in osmoregulation (32, 41), and differences in total cAMP content were observed in *TcCARP3* mutants, we next evaluated RVD in these parasites, as described in *Materials and Methods*. *TcCARP3*-KO, *TcCARP3*-AB and T7/Cas9 epimastigotes were exposed to hypoosmotic stress and the area under the curve (AUC) in different sections of the light scattering pattern was then quantified to determine the maximum volume change (AUC at 200-300 s) and the final volume recovery (AUC at 800-900 s) of the cells (Fig. 5A-C). Our results indicate that *TcCARP3*-KO parasites have a defect in their osmoregulatory capacity, as their final volume recovery was significantly higher than that of control and *TcCARP3*-AB parasites, which restored the normal phenotype of the T7/Cas9 cells (Fig. 5C). Concomitantly, *TcCARP3*-OE parasites showed a more efficient RVD profile compared to that of the EV control, exhibiting a lower maximum volume change and a more efficient final volume recovery in response to hypoosmotic stress (Fig. 5D-F). These results indicate that TcCARP3 plays an important role in the osmoregulatory capacity of *T. cruzi* epimastigotes in response to hypoosmotic conditions.

**Figure 5.**
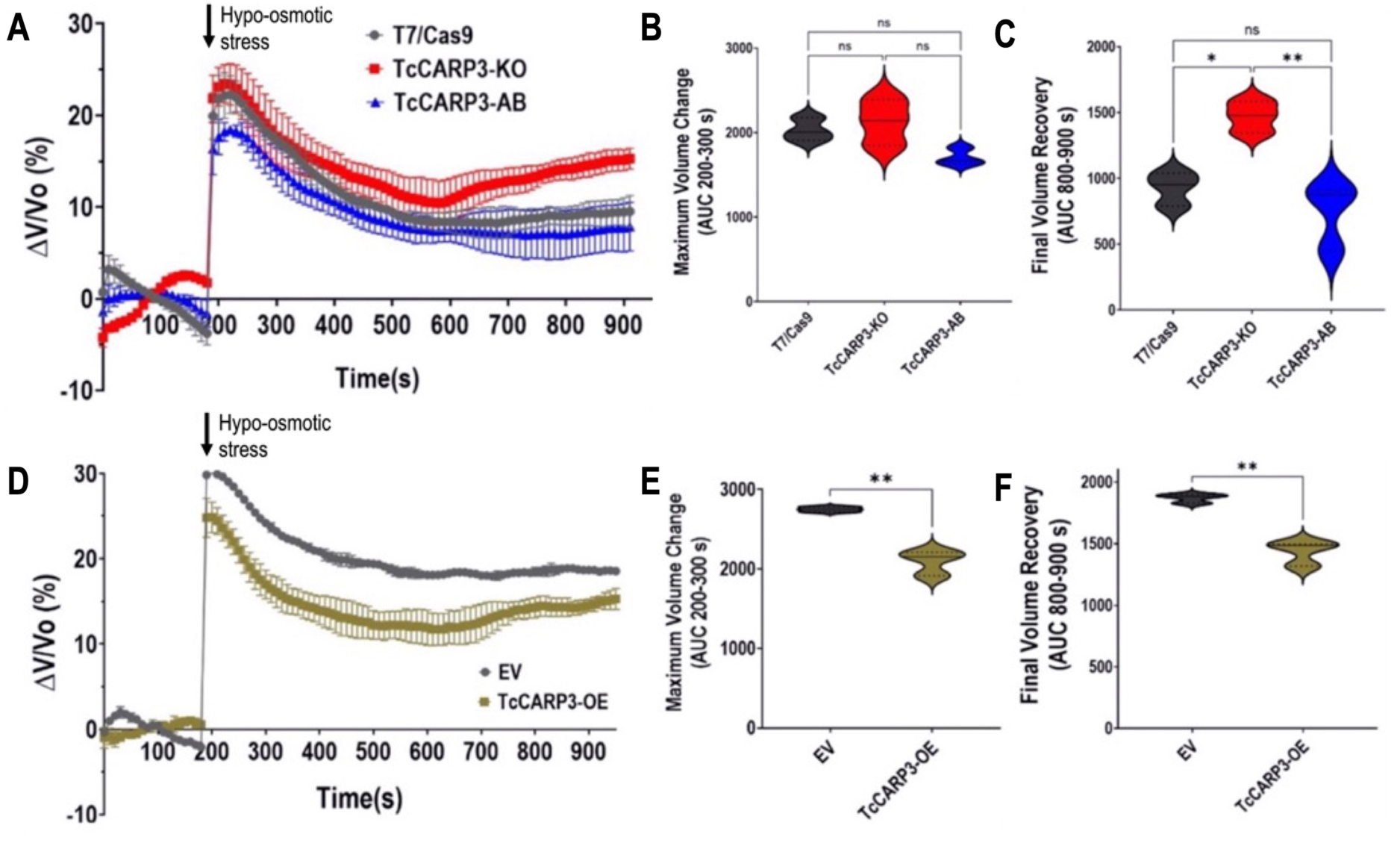
Regulatory Volume Decrease (RVD) of TcCARP3 mutants under hypoosmotic stress. **[A]** The light scattering pattern of *T. cruzi* epimastigotes suspended in isosmotic buffer was recorded for 120 s and diluted to a final osmolarity of 115 mOsm/L under constant ionic conditions. Relative changes in cell volume were monitored by measuring absorbance at 550 nm over time in T7/Cas9, *TcCARP3*-KO, *TcCARP3*-AB parasites. The absorbance values were normalized to the initial volume under isosmotic conditions and expressed as percentage of volume change. **[B]** Analysis of the maximum volume change under hypoosmotic conditions. The area under the curve (AUC) in **A** was calculated between 200 and 300 seconds for all cell lines. **[C]** Final volume recovery calculated as the AUC in **A** between 800 and 900 seconds. Values are mean ± SD; n = 3; *P < 0.05; **P < 0.01; ns, not significant differences with respect to control cells (One way ANOVA with Tukey’s multiple comparison). **[D]**, **[E]**, and **[F]** Same experiments as in **A**, **B**, and **C**, but using cell lines Empty Vector (EV) and *TcCARP3*-OE. Values are mean ± SD; n = 3. **P< 0.01 (Student’s t-test).

### Analysis of TcCARP3 protein interactors

The co-localization of TcCARP3 and TcAC1 suggested that these proteins interact in different developmental stages of *T. cruzi* life cycle, as previously observed in *T. cruzi* epimastigotes by mass spectrometry analysis (16). To confirm the interaction between TcCARP3 and TcAC1, we performed a co-immunoprecipitation assay using a dually tagged cell line (TcAC1-3xHA/TcCARP3-3xc-myc) obtained in our laboratory (16) and HA magnetic beads to trap TcAC1. Different fractions were collected and analyzed by western blot using anti-c-Myc antibodies. We were able to detect TcCARP3 in the eluted fraction (E) and the band was enriched compared to that in the third wash (3W) (Fig. 6A). Then, we co-immunoprecipitated TcCARP3 and TcAC1 using total lysates of the dually tagged cell line and c-Myc magnetic beads to trap TcCARP3. Once again, different fractions were analyzed by western blot, now using anti-HA antibodies. Likewise, we were able to detect TcAC1 in the eluate and the band was clearly enriched compared to the third wash (Fig. 6B).

**Figure 6.**
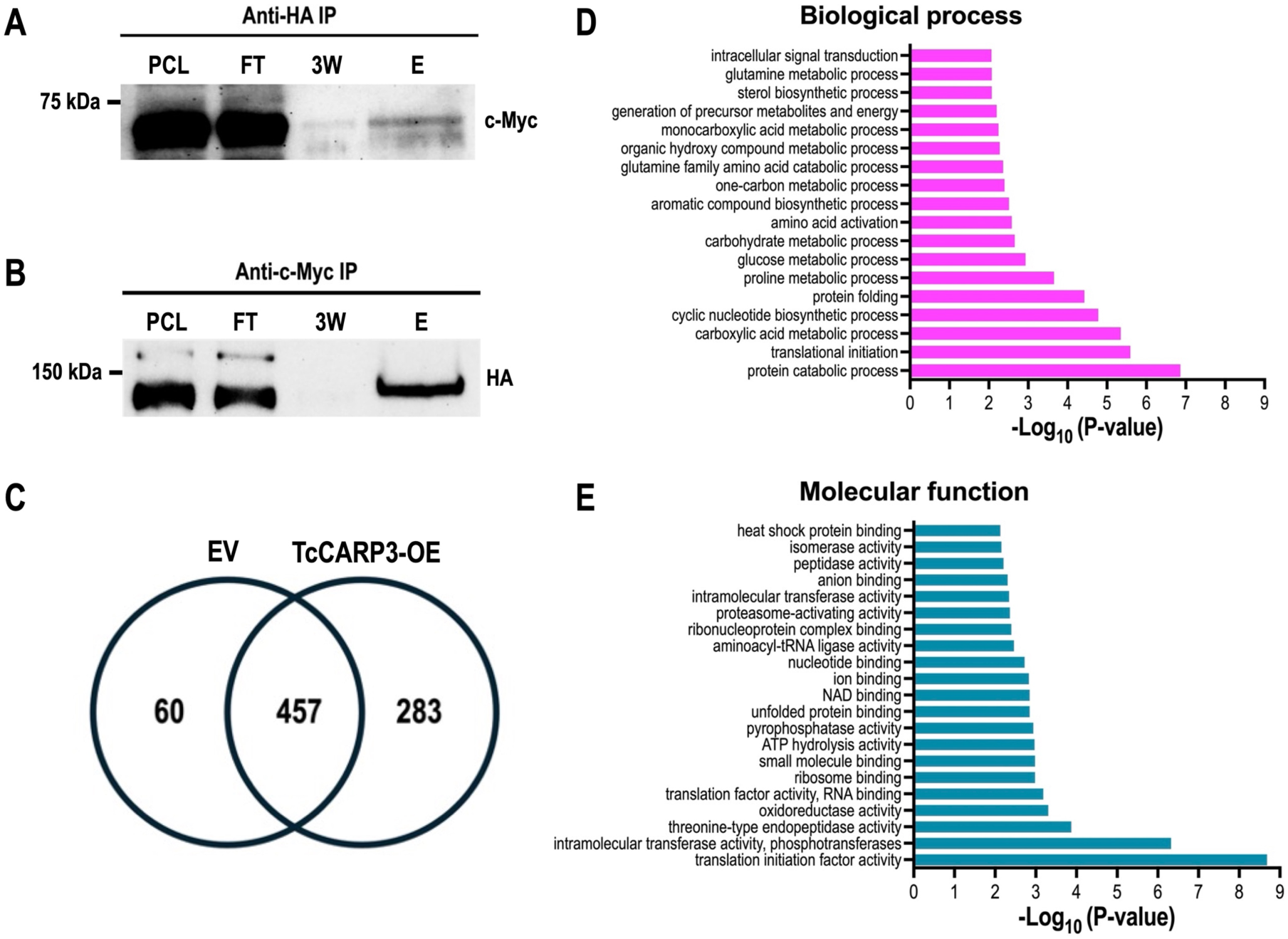
Co-IP of TcAC1-3xHA and TcCARP3-3xc-Myc, and analysis of TcCARP3 interactome. Lanes from left to right are pre-cleated lysate (PCL), flow through (FT), third wash (3W), and eluate (E). **[A]** Immunoprecipitation with HA bads to capture TcAC1 as bait followed by anti-c-Myc western blot analysis to detect TcCARP3 as prey (64.4kDa). **[B]** Immunoprecipitation with c-Myc beads to capture TcCARP3 as bait followed by anti-HA western blot analysis to detect TcAC1 as prey (144kDa). The upper band seen in the PCL and FT corresponds to Cas9-HA (>150kDa). **[C]** Venn diagram showing the number of proteins found in pTREXn-3xHA Empty Vector (EV) eluates only (60 proteins), the number of proteins found in TcCARP3-OE eluates only (283 proteins), and the number of proteins that were found in both EV and TcCARP3-OE eluates (457 proteins). **[D]** and **[E]** Gene Ontology (GO) enrichment analysis for biological process and molecular function of TcCARP3 interacting partners that were absent in the EV control (P < 0.05).

After confirming the interaction between TcCARP3 and TcAC1 we pursued to identify other TcCARP3 protein interactors. For this, we performed an immunoprecipitation assay as described above, using TcCARP3-3xHA as bait protein. In this assay, pTREXn-3xHA empty vector cell line was used as a control. After immunoprecipitation, snap frozen eluates were sent to the Mass Spectrometry Core Laboratory at the University of Texas Health Science Center at San Antonio (San Antonio, TX) for mass spectrometry analysis. The group of proteins enriched in the TcCARP3-OE cell line eluates that were absent in the EV control (infinite fold change), with a P value <0.05 after Benjamini-Hochberg multiple test correction, and a total spectra count in each TcCARP3-OE replicate ≥1, were deemed as TcCARP3 specific protein interactors (Fig. 6C, and Table S1). Interestingly, we found that TcCARP3 interacts with at least 7 adenylate cyclases, from TcAC groups I, III and V, including TcAC1 (16), and with the regulatory subunit of a PKA-like protein (PKArL), containing two putative cyclic nucleotide binding domains. We also detected several putative proteins involved in cell signaling, such as UNC119, Galactokinase-like protein, cAMP dependent protein kinase catalytic subunit 2, and a homoserine kinase, evidencing the presence of multiple signaling components sharing a subcellular niche with TcCARP3 (Table 1). A gene ontology (GO) enrichment analysis for biological process and molecular function of the 283 TcCARP3 protein interactors is shown in Figures 6D and E. Taken together, our results suggest that TcCARP3 is a multi-adenylate cyclase regulator that physically interacts with several ACs and with putative cAMP effectors in two cAMP signaling microdomains of *T. cruzi*.

**Table 1.** TcCARP3 protein interactors involved in cell signaling.

| **Product Description** | **Gene ID** | **Total spectra count** | **MW (kDa)** |
| --- | --- | --- | --- |
| Putative receptor-type adenylate cyclase (TcAC3.3) | TcYC6_0073080 | 105/87/118 | 136 |
| Putative receptor-type adenylate cyclase (TcAC3) | TcYC6_0073060 | 101/88/112 | 136 |
| Receptor-type adenylate cyclase, putative (TcAC1.1) | TcYC6_0122100 | 38/43/44 | 141 |
| Receptor-type adenylate cyclase, putative (TcAC1.5) | TcYC6_0051800 | 38/33/45 | 141 |
| Receptor-type adenylate cyclase, putative (TcAC1.4) | TcYC6_0051770 | 34/30/41 | 141 |
| UNC119 (GMP PDE delta subunit) | TcYC6_0076300 | 32/37/39 | 23 |
| Receptor-type adenylate cyclase, putative (TcAC1) | TcYC6_0015740 | 22/32/30 | 141 |
| Hypothetical protein, conserved (Protein kinase-like domain) | TcYC6_0066800 | 17/21/22 | 95 |
| Regulatory subunit of protein kinase A-like protein, putative (PKArL) | TcYC6_0089970 | 21/20/25 | 63 |
| Putative receptor-type adenylate cyclase (TcAC5.1) | TcYC6_0051420 | 20/19/29 | 134 |
| Galactokinase-like protein, putative | TcYC6_0068800 | 13/20/21 | 52 |
| cAMP-dependent protein kinase catalytic subunit 2 (PKAC2) | TcYC6_0070220 | 7/8/06 | 38 |
| Homoserine kinase | TcYC6_0098420 | 6/5/07 | 36 |

### *TcCARP3* ablation impairs the parasite’s ability to colonize the hindgut of the triatomine vector

To progress into their life cycle and undergo a successful transmission to the mammalian host, *T. cruzi* parasites must establish an efficient infection in the triatomine vector, the kissing bug, and finally reach the hindgut, where replicative epimastigotes differentiate into infective metacyclic trypomastigotes. Our results from metacyclogenesis *in vitro* indicate that TcCARP3 is involved in this differentiation process. To provide further evidence, we assessed the ability of *T. cruzi* mutants showing different expression levels of TcCARP3 to establish an infection in the triatomine bug *Rhodnius prolixus*, as described in *Materials and Methods*. Our results indicate that *TcCARP3*-KO parasites show a significantly reduced capacity to colonize the hindgut of kissing bugs, as compared to control parasites. Interestingly, these parasites were able to differentiate into metacyclic trypomastigotes. A mixed population of epimastigotes and metacyclic trypomastigotes was observed in the hindgut of infected kissing bugs, but the number of infected insects was significantly lower. The normal phenotype was rescued in the *TcCARP3*-AB cell line (Fig. 7), indicating that TcCARP3 is necessary for *T. cruzi* to establish an efficient infection in the triatomine vector. Taken together, our results shed light on the importance of TcCARP3 for the progression of *T. cruzi* life cycle *in vivo*.

**Figure 7.**
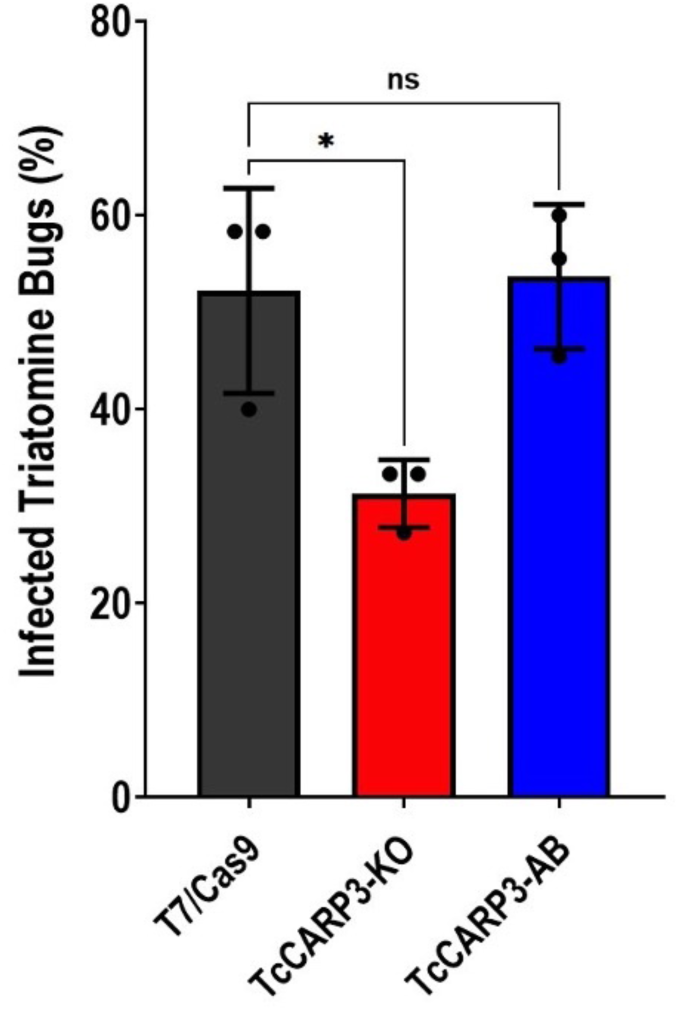
Percentage of infected triatomine bugs *in vivo*. Kissing bugs were fed blood laden with *T. cruzi* epimastigotes and hindguts were dissected 30 days later and examined on microscope for presence of parasites. *P< 0.01 (One way ANOVA with Tukey’s multiple comparisons). Values are mean ± S.D., n=3 independent experiments with 9-15 insects per group infected with each specific *T. cruzi* cell line.

## DISCUSSION

Previous results from our group showed a peculiar localization for TcCARP3 in two putative cAMP signaling microdomains: the contractile vacuole complex and the flagellar tip of *T. cruzi* epimastigotes, where this protein co-localizes with TcAC1 (16). Other cAMP signaling components have been identified in these locations: TcAC4, TcAC5, and TcPDEC2 in the CVC (15, 16), TcAC2 (CVC and flagellar tip) (16), and TcPDEB1 and TcPDEB2 along the flagellum (13, 14), supporting the idea that these two subcellular compartments are indeed cAMP signaling microdomains. We have now confirmed the co-localization of TcCARP3 and receptor-type TcAC1 in three additional developmental stages of *T. cruzi*: metacyclic trypomastigotes, amastigotes and cell-derived trypomastigotes, where their interaction modulates the levels of cAMP in this parasite. Through the generation of *TcCARP3* mutants in which this gene has been either ablated or overexpressed, we also demonstrated that TcCARP3 plays a key role in the regulation of cell volume under hypoosmotic stress and in the ability of the parasite to grow and differentiate *in vitro*, invade mammalian cells and replicate within them, as well as to colonize the digestive tract of the triatomine vector. Furthermore, we identified several adenylate cyclases and other signaling proteins as main interacting partners of TcCARP3, confirming its role as regulator of compartmentalized cAMP signals in *T. cruzi*. The interaction of CARP3 with several adenylate cyclases has been also observed in the flagellar tip of the salivarian trypanosome *T. brucei*, where this protein plays a role in SoMo (38). However, here we showed the interaction of TcCARP3 with various TcACs that localize in 2 different subcellular compartments (flagellar tip and CVC) (16), and explored new cellular processes that are modulated by this protein in *T. cruzi*. Our data highlights the relevance of TcCARP3 as regulator of compartmentalized cAMP signals throughout the life cycle of *T. cruzi*, a stercorarian trypanosome that is an obligate intracellular parasite.

The visualization of the CVC in *T. cruzi* using conventional microscopy methods is facilitated by exposing the parasites to hypoosmotic stress prior fixation (16, 34, 40). Under this condition, we observed a dual localization of TcCARP3 in epimastigotes, and in the mammalian forms of *T. cruzi*. Interestingly, we found that in metacyclic trypomastigotes TcCARP3 and TcAC1 co-localize to the flagellar distal domain, but not to the CVC under hypoosmotic conditions. Metacyclic trypomastigotes are extremely slender forms with a smaller CVC than other developmental stages (35, 53). TcCARP3 and TcAC1 may not have been detected in the CVC due to its small size in this developmental stage. Another possibility is that these proteins are not present at all in the CVC of these infective forms. The redistribution of TcCARP3 in metacyclic tryposmastigotes raises new questions about the role of the CVC in different developmental stages of *T. cruzi*.

TcCARP3 exhibits a high confidence predicted myristoylation site on the first glycine residue (second amino acid of the protein). Removal of this signal from the N-terminus of the protein did not affect the dual localization of CARP3 to the flagellar tip and the CVC in *T. cruzi* epimastigotes. This result suggests that unlike what was observed in the *T. brucei* ortholog (38), the predicted myristoylation site of TcCARP3 is either nonfunctional or it is not required for TcCARP3 localization in *T. cruzi*. However, we cannot rule out the presence of non-predicted posttranslational modifications in TcCARP3 that could determine its subcellular localization. *Trypanosoma brucei* flagellar member 8 (TbFLAM8) is necessary for TbCARP3 localization to the flagellar tip (38). A similar trafficking mechanism could be directing TcCARP3 to the flagellar tip of *T. cruzi*. However, our mass spectrometry data did not reveal TcFLAM8 as an interacting partner of TcCARP3 in *T. cruzi* epimastigotes. Further research is needed to elucidate the trafficking mechanism of TcCARP3 to these two subcellular compartments.

The flagellar distal domain is a crucial structure for *T. cruzi* attachment and subsequent metacyclogenesis in the hindgut of the triatomine bug (5, 16, 27). We previously observed an increase in cell adhesion during metacyclogenesis when the adenylate cyclase TcAC1 was overexpressed in *T. cruzi* (16). We would then expect increased cAMP local levels in the flagellar tip of *TcCARP3*-KO parasites, in which metacyclogenesis is significantly higher than in control cells. However, due to limitations of the methodology used to measure cAMP content, we were only able to estimate the relative total cAMP content in these parasites. To evaluate the specific cAMP concentration in different subcellular compartments, a biosensor cell line expressing a genetically encoded cAMP indicator should be used, as those available for mammalian cells (10). However, this technology has not been developed in trypanosomatids and so far, we cannot establish a link between local cAMP levels at the flagellar tip and the increased metacyclogenesis observed in *TcCARP3*-KO parasites, as previously reported (16, 27). Attachment of epimastigotes to the lipidic rectal cuticle precedes metacyclogenesis in the kissing bug (reviewed by (5, 39)). Interestingly, here we did not observe an adhesion defect in *TcCARP3*-KO epimastigotes during this differentiation process. Although adhesion is necessary for metacyclogenesis, it is not the only event driving this process. Indeed, there are many biochemical, morphological, and genetic changes that occur during metacyclogenesis (53). Our results suggest that the increased metacyclogenesis observed in *TcCARP3*-KO parasites is not the consequence of a cell adhesion defect. Further research should be performed to elucidate the specific role of TcCARP3 in *T. cruzi* metacyclogenesis. Analysis of gene expression profiles by single cell RNAseq at different time points during metacyclogenesis of *TcCARP3*-KO parasites could be useful to identify specific altered cellular processes in these mutants.

The second subcellular compartment where TcCARP3 localizes is the contractile vacuole complex. Among other functions, this organelle is involved in the regulation of cell volume under hypoosmotic conditions. In this process, the parasite releases water out of the cell body through an adhesion plaque in the flagellar pocket by pulsatile contractions of the central vacuole (bladder) of the CVC (35). Water efflux follows the tubulin-mediated fusion of acidocalcisomes to the central vacuole and further translocation of the aquaporin (TcAQP) to the CVC in a process known as regulatory volume decrease (32, 34, 41). RVD is a cAMP-mediated process (34, 54) that plays a key role in the survival of *T. cruzi* to extreme osmolarity fluctuations throughout its life cycle. For example, the parasite faces a dramatic drop in osmolarity when it transitions from the hindgut of the triatomine vector (∼1000mOsm/kg) to the cytosol of the mammalian host (300mOsm/kg) (7). Modulating the expression levels of TcCARP3 in *T. cruzi* caused significant changes in the ability of parasites to respond to hypoosmotic stress in their extracellular environment. In the absence of TcCARP3, *T. cruzi* epimastigotes displayed an initial swelling and subsequent volume recovery, but these parasites were not able to maintain the cell volume for more than 7 minutes (420 s) after hypoosmotic stress exposure. This initial volume recovery is compatible with the rapid release of amino acids and other inorganic osmolytes to the extracellular medium, presumably through membrane channels and transporters. This mechanism is responsible for ∼50% of cell volume recovery in *T. cruzi* (55). The remaining volume recovery is mediated by the contractile vacuole complex as described above (34, 54). Therefore, the observed *TcCARP3*-KO phenotype is indeed a defect in the CVC-mediated osmoregulatory capacity of the parasites. Conversely, when TcCARP3 is overexpressed, the parasites swell less upon hypoosmotic stress and recover better when compared to control cells. Our results support the hypothesis that abnormal levels of cAMP are generated in the CVC of these mutants. This data support a model of compartmentalized cAMP signals mediating the osmoregulatory capacity of *T. cruzi*.

To gain further insight into how TcCARP3 might differentially modulate cAMP synthesis at the flagellar tip and contractile vacuole complex, we analyzed the interactome of TcCARP3 on *T. cruzi* epimastigotes. We previously demonstrated the physical interaction of TcCARP3 and TcAC1 by immunoprecipitation assays and mass spectrometry analysis (16). Here, we confirmed this interaction through co-IP assays and mass spectrometry analysis using TcCARP3 as bait. Mass spectrometry revealed that TcCARP3 not only interacts with TcAC1 but also with other seven adenylate cyclases from three different groups (TcAC groups I, III and V). TcAC3 (group III) was previously observed in the ER of epimastigotes, as shown by partial co-localization with the ER marker BiP (16). Considering this new evidence, the localization of TcAC3 should be further investigated to determine if it is indeed localized to the CVC with accumulation in the ER due to overexpression. The *T. brucei* irtholog TbCARP3 was also found to interact with multiple adenylate cyclases in *T. brucei*, and to modulate them in different ways depending on the specific identity of the interactor (38). How TcCARP3 interacts with different TcACs and the specific downstream effects arising from these interactions are questions that should be further investigated. In this regard, the predicted TPR-like tetratricopeptide-like helical domain in TcCARP3 (InterPro ID: IPR011990) could be mediating protein-protein interactions (50). Interestingly, this domain is not predicted by InterProScan (56) in the *T. brucei* homolog TbCARP3.

Ablation of *TcCARP3* leads to phenotypes compatible with increased cAMP levels in one microdomain (the flagellar tip), and decreased cAMP content in another microdomain (the CVC). Furthermore, the abundance of TcACs detected by mass spectrometry differs between groups (AC3 > AC1 > AC5 isoforms), suggesting that TcCARP3 shows specific affinities for different AC groups. Since trypanosome ACs become catalytically active upon dimerization (57), the monomers to dimers proportion and their composition (homo or heterodimers) could be modulated by TcCARP3-TcAC interactions, determining the TcAC catalytic state and cAMP content in these specific microdomains. We hypothesize that trypanosome AC dimerization state is the indirect consequence of membrane modifications occurring in response to microenvironmental cues, which affect AC-CARP3 interactions in membrane microdomains. Taken together, our results suggest that TcCARP3 is a multi-adenylate cyclase regulator, where the TcAC identity may determine the catalytic state of different AC dimers. In addition to interacting with several TcACs, our mass spectrometry data indicates that TcCARP3 interacts with the regulatory subunit of a PKA-like protein, a protein that contains two putative cyclic nucleotide binding domains, and therefore could be a cAMP effector. PKArL is the homolog of a divergent protein kinase A regulatory subunit (PKAR3) recently described in *Leishmania donovani* (58). This protein is absent in most trypanosomatids, including *T. brucei*, and is essential for maintenance of the elongated shape of *Leishmania* promastigotes. The interaction of TcCARP3 with PKArL provides further evidence on the role of TcCARP3 in the cAMP signal transduction pathway in *T.* cruzi.

The role of cAMP in osmoregulation and metacyclogenesis has been reported by several groups over the last decades (7, 15, 16, 27, 29-34, 54). However, its role in host cell invasion and intracellular replication was first described by our laboratory (16). During the characterization of TcAC1, we showed that increased levels of cAMP leads to a defect in the ability of *T. cruzi* trypomastigotes to invade mammalian host cells and replicate intracellularly as amastigotes. A similar defect was observed in *TcCARP3*-KO parasites showing a lower number of infected host cells at 24 h post-infection, and a lower number of intracellular amastigotes after 72 h in mammalian cell cultures. These results further demonstrate that cAMP plays a key role in environmental sensing, with implications for life cycle progression of *T. cruzi* in the mammalian host, as TcCARP3 mutant parasites exhibit decreased levels of cAMP. It would be interesting to evaluate if these mutants are able to establish an acute infection in a murine model. Interestingly, *TcCARP3*-KO parasites showed a defect in colonizing the digestive tract of kissing bugs. Besides their metacyclogenesis phenotype displayed *in vitro*, these parasites were not able to efficiently establish an infection in the insect vector. A possible explanation is that cells undergoing *in vitro* metacyclogenesis in TAU3AAG are subjected to conditions mimicking the vector’s urine composition, but not to the osmolarity the parasite is exposed in the triatomine bug. These conditions (nutrient deprivation and low pH) (59), trigger differentiation from replicative epimastigotes to infective metacyclic trypomastigotes. However, in the hindgut of the kissing bug, where metacyclogenesis naturally occurs, the microenvironment within the vector reaches an extremely high osmolarity of ∼1000 mOsm/kg (7) while in TAU3AAG medium the osmolarity is about ∼300 mOsm/kg. It is possible that *TcCARP3*-KO parasites do not tolerate the osmotic stress it faces in the vector’s gastrointestinal tract, since these parasites showed a reduced osmoregulatory capacity. While the absence of TbCARP3 in the salivarian *T. brucei* was also found to hinder the colonization of tissues in its arthropod vector, the tsetse fly (37, 38), this probably occurs in response to specific microenvironmental cues that are different to those faced by *T. cruzi* in the triatomine vector. Ablation of TbCARP3 did not affect the parasite’s ability to differentiate from procyclic forms to epimastigotes and then to metacyclic trypomastigotes *in vitro*, but caused a defect in social motility, and in the ability of parasites to colonize the tsetse fly salivary glands (37, 38). *T. cruzi* on the other hand does not encounter physical barriers in the triatomine vector but does experience intense osmotic stress to which *T. brucei* is never exposed to during its life cycle. In this regard, osmotic stress could have been a driving force in the evolutive retention of the CVC in *T. cruzi,* and for the development of a cAMP signaling microdomain in this subcellular compartment. Importantly, our kissing bug infection results represent the first report of loss-of-function analysis using *T. cruzi* mutant cell lines to infect the triatomine vector.

Mounting evidence on the role of cAMP in environmental sensing in trypanosomes and in other protozoan parasites has been reported during the last 20 years (7, 15, 16, 19, 27, 33, 34, 37, 42, 60-62). Our data demonstrates that TcCARP3 modulates cAMP levels in *T. cruzi*, and is involved in osmoregulation, metacyclogenesis, host cell invasion, intracellular replication and colonization of the vector’s digestive tract, providing relevant new evidence on the role of cAMP in environmental sensing. We also found that TcCARP3 co-localizes with TcAC1 in all four developmental stages of *T. cruzi* and further demonstrated direct interaction between these proteins. Together with our proteomic data, these results significantly add to the body of evidence supporting that TcCARP3 is a multi-adenylate cyclase regulator in the flagellar distal domain and the contractile vacuole complex of *T. cruzi*. Future research should be oriented to elucidate the nature of TcCARP3 protein interactions, specifically with adenylate cyclases and putative cAMP effectors, and to determine how TcCARP3 modulates their activity. Characterizing the signaling components in individual cAMP microdomains is crucial to unveil the essential regulatory mechanisms driving cAMP signaling in trypanosomes.

## MATERIALS AND METHODS

### Chemicals and reagents

Fetal bovine serum (FBS) was purchased from R&D Systems (Minneapolis, MN). G418 was obtained from KSE Scientific (Durham, NC). Puromycin, blasticidin S HCl, Subcloning Efficiency DH5a competent cells, BCA Protein Assay Kit, SuperSignal West Pico Chemiluminescent Substrate, Horseradish peroxidase (HRP)-conjugated anti-mouse and anti-rabbit IgG antibodies, mouse anti-HA monoclonal antibody, and Pierce™ Anti-HA and Anti-c-Myc Magnetic Beads were from Thermo Fisher Scientific (Waltham, MA). Alexa Fluor 488-conjugated donkey anti-mouse, and Alexa Fluor 594-conjugated donkey anti-rabbit were from Jackson ImmunoResearch (West Grove, PA). Restriction enzymes, and Q5^®^ High-Fidelity DNA Polymerase were obtained from New England BioLabs (Ipsich, MA). ZymoPURE Plasmid Miniprep, ZymoPURE II Plasmid Midiprep and DNA Clean & Concentrator-5 were from Zymo Research (Irvine, CA). cAMP-Glo™ Assay kit, T4 DNA Ligase and GoTaq G2 Flexi DNA Polymerase were from Promega (Madison, WI). cOmplete™ Mini EDTA-free Protease Inhibitor Cocktail was from Roche (Basel, Switzerland). 4-mm electroporation cuvettes, Precision Plus Protein Dual Color Standards, and nitrocellulose membranes were from Bio-Rad (Hercules, CA). Mouse anti-c-Myc monoclonal antibody (9E10) was from Santa Cruz Biotechnology (Dallas, TX). Fluoromount-G mounting medium was from Southern Biotech (Birmingham, AL). The pMOTag23M vector (63) was a gift from Dr. Thomas Seebeck (University of Bern, Bern, Switzerland). DNA oligonucleotides were purchased from Integrated DNA Technologies (Coralville, IA). Phenylmethylsulfonyl fluoride (PMSF), N-*p*-tosyl-L-phenylalanine chloromethyl ketone (TPCK), trans-epoxysuccinyl-l-leucylamido-(4-guanidino)butane (E64), protease inhibitor cocktail for use with mammalian cell and tissue extracts (Cat. No. P8340), Benzonase nuclease, and all other reagents of analytical grade were from Sigma-Aldrich (St. Louis, MO). Adult *Rhodnius prolixus*, Strain CDC, NR-44077 was provided by Centers for Disease Control and Prevention for distribution by BEI Resources, NIAID, NIH.

### In silico analyses

*TcCARP3* (TriTrypDB gene ID: TcYC6_0045920) sequence and reported tetratricopeptide-like helical domain were retrieved from TriTrypDB.org (44). Proteolipid modification predictions such as myristoylation prediction were done using the Research Institute of Molecular Pathology (IMP) NMT – The MYR Predictor (45) and the GPS-Lipid (lipid.biocuckoo.org) tools. Sequence alignment of nucleotides and amino acids was performed using VectorBuilder (vectorbuilder.com). *In silico* restriction enzyme digests, primer designs, and Alpha Fold 3D structure predictions were carried out using Benchling.

### Cell cultures

*T. cruzi* epimastigotes (Y strain) were grown in culture flasks containing liver infusion tryptose (LIT) medium (64) supplemented with 10% heat-inactivated fetal bovine serum (FBS), penicillin (100 I.U./mL), and streptomycin (100 µg/mL) at 28°C. Cell density was determined using a Neubauer hemocytometer counting chamber. Control parasites transfected with pTREX-n-3xHA empty vector, and overexpressing cell lines of *TcAC1*-OE and *TcCARP3*-OE were grown in the presence of 250 µg/mL G418. *TcCARP3*-3xc-Myc and *TcCARP3*-3x-Ty1 endogenously tagged cell lines were maintained with 250 µg/mL G418 and 5 µg/mL puromycin. Dually tagged *TcCARP3*-3x-c-Myc/*TcAC1*-3xHA and *TcCARP3*-KO cell lines were grown with 250 µg/mL G418, 5 µg/mL puromycin and 10 µg/mL blasticidin. *TcCARP3*-AB and *TcCARP3*-8AA were grown in 250 µg/mL G418, 5 µg/mL puromycin, 10 µg/mL blasticidin, and 250 µg/mL Hygromycin. Tissue culture-derived trypomastigotes and amastigotes were collected from the culture medium of infected human foreskin fibroblasts (hFFs) cells. hFFs were grown in DMEM (Dulbecco’s Modified Eagle Medium, Gibco) supplemented with 10% FBS, penicillin (100 I.U./mL), and streptomycin (100 µg/mL), and maintained with 5% CO_2_ at 37°C.

### Generation of *TcCARP3* overexpression and dually tagged *TcCARP3*/*TcAC1* parasites

The open reading frame of *TcCARP3* was PCR amplified using *T. cruzi* Y strain gDNA as template (primers 1 & 2; Table S2) and cloned into pTREX-n-3xHA vector (65) by restriction sites XbaI/EcoRV. The construct pTREX-b-*TcAC1*-3xHA for the dually tagged cell line of *TcAC1*-3xHA/*TcCARP3*-3xc-Myc was generated as described in (16). Briefly, amplification of *TcAC1*-3xHA was done by PCR using pTREX-n-*TcAC1*-3xHA plasmid as template and then cloned into pTREX-b by HindIII restriction site using NEBuilder® HiFi DNA Assembly cloning kit (New England Biolabs). Gene cloning was confirmed by sequencing, and constructs were used to transfect *T. cruzi* epimastigotes. The pTREXn-*TcCARP3*-3xHA construct was used to transfect WT cells to obtain an overexpression cell line *TcCARP3*-OE Clonal populations were obtained by serial dilutions. The pTREXb-*TcAC1*-3xHA construct was used to transfect endogenously tagged *TcCARP3*-3xc-Myc epimastigotes to obtain a dually tagged cell line. Expression of TcCARP3 and TcAC1 was confirmed by western blot analysis using anti-c-Myc and anti-HA antibodies, respectively.

### Ablation of *TcCARP3*

We performed a CRISPR/Cas9-mediated knock out of *TcCARP3* using a standard strategy developed in our laboratory (66). Briefly, *Trypanosoma cruzi* Y strain epimastigotes constitutively expressing T7 RNA polymerase and Cas9 were transfected with a sgRNA template obtained by PCR (primers 3 & 17; Table S2) and two donor DNA cassettes amplified from pGEM-BSD-TGA and pGEM-PAC-TGA (primers 4 & 5; Table S2), respectively. The donor DNA was provided to induce homology-directed repair (HDR) and contained a blasticidin or a puromycin resistance marker respectively flanked by 40 and 37-nt homologous regions corresponding to the 5’ and 3’ end of the *TcCARP3* UTRs. Selection of the protospacer was performed using EuPaGDT (eukaryotic pathogen CRISPR guide RNA/DNA design tool; http://grna.ctegd.uga.edu) (58). We chose a specific sgRNA sequence targeting a site within the ORF of the *TcCARP3* gene. Selection of transfectants was done with puromycin and blasticidin to ensure both alleles were replaced by resistance markers. Gene knockout was verified by PCR from gDNA using a specific set of primers (primers 6 & 7; Table S2). After *TcCARP3* knockout was confirmed, a clonal population was obtained by serial dilutions.

### Generation of addback cell lines

We obtained the addback cell line by amplifying the ORF of *TcCARP3* using pTREXn-*TcCARP3*-3xHA as a template and subcloning into pTREXh-2xTy1 through restriction sites XbaI/EcoRV (primers 1 & 8; Table S2). This construct was then used to transfect *TcCARP3*-KO parasites to obtain the *TcCARP3* addback (*TcCARP3*-AB). We obtained another construct with a truncated version of TcCARP3 where the first 8 amino acids had been deleted to get rid of the first 3 glycine residues that contained a predicted myristoylation site. To achieve this, the sequence downstream of the 8^th^ codon (24 nt) of *TcCARP3* was amplified using pTREXn-*TcCARP3*-3xHA as a template and subcloned into pTREXh-2xTy1 through restriction sites XbaI/EcoRV (primers 8 & 9; Table S2). The construct was then transfected into *TcCARP3*-KO parasites to obtain the *TcCARP3*-8AA cell line. Both constructs were verified by restriction digestion and Sanger sequencing before transfection. Successfully transfected parasites were confirmed by western blot and IFA after selection with hygromycin.

### Endogenous tagging of *TcCARP3*

We performed a CRISPR/Cas9-mediated endogenous C-terminal tagging of *TcCARP3*. Briefly, *Trypanosoma cruzi* Y strain epimastigotes constitutively expressing T7 RNA polymerase and Cas9 were transfected with a sgRNA template obtained by PCR (primers 10 & 17; Table S2) and a donor DNA cassette, amplified from pMOTag23T vector (primers 11 & 12; Table S2). The pMOTag23T vector was made by amplifying two copies of the Ty1 tag from the pTREXh-2xTy1 vector (primer 13 & 14; Table S2) and cloned into pMOTag2T already containing a puromycin resistance marker and one copy of the Ty1 tag (63) by XhoI site using NEBuilder® HiFi DNA Assembly cloning kit (New England Biolabs). The Donor DNA provided to induce homology-directed repair contained a 3x-Ty1 tag, a puromycin resistance marker, and 65 and 60-nt homologous regions at the 5’and 3’ ends of the cassette, respectively. Selection of the protospacer was performed using EuPaGDT. We chose a specific sgRNA sequence targeting the 3’end of *TcCARP3 gene*. Selection of transfectants was done with puromycin. Endogenous gene tagging was verified by PCR from gDNA using a specific set of primers (primers 15 &16; Table S2) and by western blot analysis.

### Transfection of *T. cruzi* epimastigotes

*Trypanosoma cruzi* Y strain epimastigotes were transfected via electroporation as previously described (65). Briefly, 4 × 10^7^ cells in early exponential phase were washed with sterile 1x PBS pH 7.4 at RT and resuspended in ice-cold CytoMix (120 mM KCl, 0.15 mM CaCl_2_, 10 mM K_2_HPO_4_, 25 mM HEPES, 2 mM EDTA, 5 mM MgCl_2_, pH 7.6) to a final density of 1 × 10^8^ cells/mL. Then, 400 μL of cell suspension were transferred to a cold 4-mm electroporation cuvette that was on ice containing 25 μg of each DNA fragment (purified plasmid or PCR product) in a maximum DNA volume of 40 uL. Three electric pulses (1500 V, 25 μF) were sent to the cells in cuvettes, using a Gene Pulser Xcell Electroporation System (Bio-Rad). Transfected epimastigotes were cultured in LIT medium supplemented with 20% heat-inactivated FBS and the corresponding antibiotics for selection of successfully transfected parasites expressing antibiotic resistance, until healthy cell lines were obtained (2-3 weeks). Clonal populations of transfectant parasites were obtained by serial dilutions in LIT medium and a final dilution in conditioned media (20% heat inactivated FBS, 40% filtered supernatant from WT cells in exponential phase, 40% LIT media, penicillin (100 I.U./mL), streptomycin (100 µg/mL), and appropriate antibiotics) to a final density of 2.5 cells/mL and plated 200µl per well in 96-well plates.

### Western blot analyses

Western blots were performed as previously described (67). Briefly, parasites in exponential phase of growth were washed in 1x PBS pH 7.4 and resuspended in radio-immunoprecipitation assay (RIPA) buffer (150 mM NaCl, 20 mM Tris-HCl, pH 7.5, 1 mM EDTA, 1% SDS, 0.1% Triton X-100) plus a mammalian cell protease inhibitor cocktail (diluted 1:250), 1 mM phenylmethylsulfonyl fluoride, 2.5 mM tosyl phenylalanyl chloromethyl ketone, 100 M *N*- (*trans*-epoxysuccinyl)-L-leucine 4-guanidinobutylamide (E64), and Benzonase Nuclease (25 U/mL culture). After lysis the cells were then incubated for 30 min on ice, and protein concentration was determined by BCA protein assay. Thirty micrograms of protein from each cell lysate were mixed with 4x Laemmli sample buffer (Bio-Rad) supplemented with 10% β-mercaptoethanol, before loading into a 10%, 8%, or 6% SDS–polyacrylamide gels. Electrophoresed proteins were then transferred onto nitrocellulose membranes with a Trans-Blot Turbo Transfer System (Bio-Rad). After transfer the membranes were stained with Ponceau red and an image was acquired for loading control using a ChemiDoc Imaging System (Bio-Rad). Membranes were then destained using PBS-T (PBS containing 0.1% Tween 20) and blocked with 5% nonfat dry milk in PBS-T overnight at 4°C. Then, the nitrocellulose membranes were incubated for 1 hour at room temperature, with the primary antibody: monoclonal anti-HA (1:2000), monoclonal anti-c-Myc (1:1000), or monoclonal anti-Ty1 (1:2000). After three washes with PBS-T, blots were incubated with the secondary HRP-conjugated antibody (goat anti-mouse IgG or goat anti-rabbit IgG, diluted 1:10,000). Membranes were washed three times with PBS-T and incubated with Pierce™ ECL Western Blotting Substrate (Thermo Fisher Scientific) in dark for 5 min. Lastly, images were acquired with a ChemiDoc Imaging System (Bio-Rad).

### Immunofluorescence analyses

*T. cruzi* parasites (epimastigotes, trypomastigotes or amastigotes) were washed with 1x PBS pH 7.4 and fixed with 4% paraformaldehyde (PFA) in 1x PBS pH 7.4 for 1 h at RT. IFAs involving TcAC1 and TcCARP3 mutants were performed under hypoosmotic conditions, by adding and equal volume of deionized water to the parasites in 1x PBS pH 7.4 and fixing them after exactly 2 min. Thereafter, cells were allowed to adhere to 1 mg/mL poly-L-lysine–coated coverslips and then permeabilized for 5 min with 0.1% Triton X-100. The coverslips were then washed 3 times with 1x PBS pH 7.4. Cells were then blocked with trypanosome blocking solution (3% bovine serum albumin (BSA), 1% fish gelatin, 5% normal goat serum and 50 mM NH_4_Cl, in PBS pH 7.4), overnight at 4°C. Next the cells were incubated with primary antibodies: rabbit anti-HA (1:200) and/or mouse anti-c-Myc (1:100), diluted in 1% BSA in 1x PBS pH 8.0 for 1 h at RT. Cells were washed three times with 1% BSA in 1x PBS pH 8.0 and then incubated for 1 h at RT with secondary antibodies: Alexa Fluor 488–conjugated donkey-anti mouse (1:400) and/or Alexa Fluor 594–conjugated donkey-anti rabbit (1:400). The incubation was performed keeping the cells protected from light to avoid photobleaching. Then, cells were washed 3 times with 1% BSA in 1x PBS pH 8.0 and mounted on slides using Fluoromount-G mounting medium containing 5 μg/mL 4,6-diamidino-2-phenylindole (DAPI) to stain genetic material. Differential interference contrast (DIC) and fluorescence optical images were captured using a Nikon Ni-E epifluorescence microscope on 100x oil immersion lens using NIS-Elements software for acquisition and subsequent processing of the images.

### Determination of cAMP content

Intracellular levels of cAMP in *T. cruzi* epimastigotes were determined using the luminescent assay cAMP-Glo™ (Promega) following manufacturer’s protocol. Briefly, *T. cruzi* epimastigotes in exponential phase of growth were washed twice with 1x PBS pH 7.4 and resuspended in induction buffer (500 mM 3-Isobutyl-1-methylxanthine and 100 mM Ro 20-1724 in PBS, pH 7.4) to a final density of 1 × 10^9^ cells/mL. Next, 10 mL of cell suspension were transferred into a white 96-well plate in triplicates (1× 10^7^ cells/well). A portion of the total lysate was used to quantify protein concentration using BCA assay. Cells in wells were lysed adding 10 mL of cAMP-Glo™ lysis buffer and incubating them at RT for 15 min. Next, 20 uL of cAMP detection solution were added to each well. Cells in plate were agitated for 1 min in an orbital shaker and incubated for 20 min at RT. Finally, 40 uL of Kinase-Glo^©^ Reagent were simultaneously added to the wells. After shaking for 1 min, the plate was incubated for 10 min at RT. Luminescence was measured using a BioTek Synergy H1 plate reader (Agilent Technologies). Results were expressed as mean values of cAMP content relative to control cells from three independent experiments and normalized by protein concentration.

### RVD assays

Regulatory volume decrease after hypoosmotic stress was monitored as described previously (16, 40). Briefly, *T. cruzi* epimastigotes in exponential phase of growth were centrifuged at 1,000 × g for 7 min, washed two times in 1x PBS pH 7.4, and resuspended in isosmotic buffer (64 mM NaCl, 4 mM KCl, 1.8 mM CaCl_2_, 0.53 mM MgCl_2_, 5.5 mM glucose, 150 mM D-mannitol, 5 mM HEPES-Na, pH 7.4, 282 mOsmol/L) at a cell density of 1×10^8^ cells/mL. Next, 100 μL was aliquoted in a 96-well plate in triplicates and the absorbance at 550 nm was measured every 10 s for 3 min using a BioTek Synergy H1 plate reader (Agilent Technologies). Then, 200 μL of hypoosmotic buffer (64 mM NaCl, 4 mM KCl, 1.8 mM CaCl_2_, 0.53 mM MgCl_2_, 5.5 mM glucose, and 5 mM HEPES-Na, pH 7.4), were added simultaneously for a final osmolarity of 115 mOsmol/L, and the absorbance at 550 nm was measured after hypoosmotic stress for additional 12 min. Readings were normalized using the mean value of the initial 3 min in isosmotic buffer. Normalized 550nm absorbance readings were then converted into a percent volume change using the equation: (|Vf – Vo| / Vo) × 100, where Vf is the absorbance value at the time point after hypoosmotic stress and Vo is the absorbance mean value obtained under isosmotic conditions. The osmoregulatory capacity of *T. cruzi* cell lines were quantified using two different parameters: the maximum change of cell volume upon induction of hypoosmotic stress (area under the curve between 200 and 300 s in the absorbance chart) and the final volume recovery (area under the curve between 700 and 800 s).

### *In vitro* metacyclogenesis

Metacyclic trypomastigotes were obtained following the protocol described in (59) with some modifications. Briefly, *T. cruzi* epimastigotes were cultured for 4 days in LIT medium supplemented with 10% heat inactivated FBS. The parasites were then washed two times in 2 mL of triatome artificial urine (TAU) (190 mM NaCl, 17 mM KCl, 2 mM MgCl_2_, 2 mM CaCl_2_, 0.035% sodium bicarbonate, 8 mM phosphate, pH 6.9) and resuspended in 0.2 mL of TAU medium. Parasites were then incubated for 2 h at 28°C. After incubation, parasites were added to flasks and incubated horizontally for 96 h in 20 mL TAU 3AAG medium (TAU medium supplemented with 10 mM L-proline, 50 mM sodium L-glutamate, 2 mM sodium L-aspartate, and 10 mM glucose) in T25 flasks. For quantification of metacyclogenesis, the supernatant containing a mixture of epimastigotes, metacyclic trypomastigotes, and intermediate forms were centrifuged at 1300 x g for 15 min and fixed for 1 h at RT in 4% PFA in PBS, attached to poly-L-Lysine-coated coverslips and washed three times with 1x PBS pH 7.4. Then, parasites were mounted onto glass slides with Fluoromount-G containing 15 μg/mL DAPI, for DNA staining. 20 fields/slide were analyzed on a Nikon epifluorescence microscope with a 100x objective under oil immersion in three independent experiments. Metacyclic trypomastigotes were distinguished from epimastigotes by the kinetoplast location in the cell body. The kinetoplast is more posterior in metacyclic trypomastigotes while in epimastigotes it is located between the nucleus and the flagellum. After confirmation of the kinetoplast location, DIC was used to confirm the slender morphology consistent with a metacyclic trypomastigote.

### Adhesion assays

During *in vitro* metacyclogenesis, parasites adhere to the plastic within the first 6 h of horizontal incubation in TAU 3AAG medium. Thereafter, fully differentiated metacyclic trypomastigotes spontaneously detach and are released into the TAU 3AAG during the following 96 h (27). To assess the ability of *T. cruzi* epimastigotes to adhere to the plastic in a sterile 12 well plate during the incubation in TAU 3AAG medium, parasite density in the medium was determined at 2, 4, 6, 24, 48, 72 and 96 h using a Neubauer hemocytometer counting chamber. The number of parasites that were adhered was determined by subtracting the total number of non-adhered cells using the density calculated and total volume added from the initial number of cells added to the well.

### Host cell invasion and intracellular replication

*T. cruzi* invasion and intracellular replication assays were performed using human foreskin fibroblasts (hFFs). First, 5×10^4^ hFFs in 1 mL of DMEM supplemented with 10% FBS were added to 12-well plates containing a sterile coverslip and allowed to attach overnight at 37°C with 5% CO_2_. The next day a swimming protocol was performed on tissue culture-derived trypomastigotes by centrifuging at 1700 x g for 15 min and incubating upright in a 50 mL conical tube for 4h at 37°C with 5% CO_2_. This allowed the competent trypomastigotes to swim out of the pellet into the supernatant. Next, the supernatant was spun down, and density of parasites was determined using a Neubauer chamber and resuspended to a concentration of 5×10^6^ parasites/mL. The hFFs in the 12-well plate were washed with DHANKS (Hank’s Balanced Salt Solution, Cytiva Marlborough, MA) and 1 mL of the parasite suspension (5×10^6^ parasites) was added for a multiplicity of infection (MOI) of 100 (100 trypomastigotes/cell). The infection was stopped after 4 h by washing the coverslips in the wells 5 times with DHANKS. Then 1 mL of DMEM with 2% FBS was added to slow down the proliferation of hFFs. Coverslips were removed from the plate after 24 h for the invasion assay, and after 72 h for the replication assay, and placed into a 12-well plate containing 4% paraformaldehyde in 1x PBS pH 7.4 for 1 h. The coverslips were then washed with 1x PBS pH 7.4 and mounted onto glass slides containing 15 μg/mL DAPI in Fluoromount G for DNA staining of parasites and mammalian cells. To quantify invasion, 20 fields/slide were visualized on a Nikon Ni-E epifluorescence microscope, and the number of infected and non-infected cells were counted. To quantify the replication of amastigotes, 60 infected host cells were visualized per assay on a Nikon Ni-E epifluorescence microscope and the number of amastigotes per infected cell were counted.

### Co-immunoprecipitation of TcAC1-3xHA/TcCARP3-3xc-Myc

*Trypanosoma cruzi* epimastigotes (2 × 10^8^ cells) in exponential phase of growth were centrifuged at 1,000 × g for 15 min and washed twice with 5 mL of buffer A with glucose (BAG, 116 mM NaCl, 5.4 mM KCl, 0.8 mM MgSO_4_, 50 mM HEPES and 5.5 mM glucose, pH 7.3) at room temperature. The parasites were then resuspended in 1 mL of ice-cold lysis buffer (0.4% NP-40, 1 mM EDTA, 150 mM KCl, cOmplete™ Mini EDTA-free Protease Inhibitor Cocktail, 50 mM Tris-HCl, pH 7.5) and mixed at 4°C for 30 min with agitation on a rocking shaker. After lysis, the protein concentration of each sample was determined using BCA assay. Cell lysate was centrifuged at 4°C at 15,000 × g for 20 min and the supernatant was incubated with 50 μL of Pierce™ Anti-HA magnetic beads to trap TcAC1-3xHA as the bait protein, or Pierce™ Anti-c-Myc magnetic beads to trap TcCARP3-3xc-myc as the bait protein. The beads had been previously washed with lysis buffer using a magnetic rack and the amount of protein loaded into the tubes with the magnetic beads was standardized based on BCA protein quantification. A portion of the pre-cleared lysate (PCL) was saved for subsequent western blot analysis. The soluble fraction of the supernatant was then incubated with magnetic beads for 1 h at RT with gentle agitation. After incubation, the flow-through was removed and saved and the magnetic beads were then washed 3 times with wash buffer (0.1% NP-40, 1 mM EDTA, 150 mM KCl, cOmplete™ Mini EDTA-free Protease Inhibitor Cocktail, 50 mM Tris-HCl, pH 7.5) using a magnetic rack. The third wash was saved for subsequent western blot analysis. A final wash was performed using pure deionized water. Proteins were then eluted with 100 μL of elution buffer (0.1M glycine, pH 2.0), by applying gentle agitation for 10 min at RT. Eluates were then neutralized with 15 μL neutralization buffer (1M Tris, pH 9.5) and analyzed by western blot with anti-HA antibodies for the anti-c-Myc IP to detect TcAC1-3x-HA as prey, or with anti-c-Myc antibodies for the anti-HA IP to detect TcCARP3-3x-c-Myc as prey.

### Analysis of TcCARP3 Interactome

We performed immunoprecipitation of TcCARP3-3xHA overexpression and pTREXn-3xHA empty vector cell lines using Pierce™ anti-HA magnetic beads. The same general procedure was followed as described above in the “Co-immunoprecipitation of TcAC1-3xHA/TcCARP3-3xc-Myc" methods section. Eluted fractions from TcCARP3-3xHA overexpressing parasites and empty vector control cells were sent to the Mass Spectrometry Core Laboratory at The University of Texas Health Science Center (San Antonio, TX) for analysis. Aliquots of the eluates (100 µL) were mixed with 100 µL 10% SDS in 50 mM triethylammonium bicarbonate (TEAB), reduced with tris(2-carboxyethyl)phosphine hydrochloride (TCEP), alkylated in the dark with iodoacetamide and applied to S-Traps micro (ProtiFi) for tryptic digestion (sequencing grade; Promega) for 2 hr in 50 mM TEAB. Peptides were eluted from each S-Trap with 0.2% formic acid in 50% aqueous acetonitrile. Digests were analyzed by capillary HPLC-electrospray ionization tandem mass spectrometry on a Thermo Scientific Orbitrap Fusion Lumos mass spectrometer. On-line HPLC separation was accomplished with an RSLC NANO HPLC system (Thermo Scientific/Dionex) interfaced with a Nanospray Flex ion source (Thermo Scientific) fitted with a PepSep column (Bruker; ReproSil C18, 15 cm x 150 µm, 1.9 µm beads). Precursor ions were acquired in the orbitrap in centroid mode at 120,000 resolution (*m/z* 200); data-dependent higher-energy collisional dissociation (HCD) spectra were acquired at the same time in the linear trap using the “rapid" speed option (30% normalized collision energy). Mascot (v2.8.3; Matrix Science, London UK) was used to search the spectra against a combination of the following databases: TcruziYC6_TriTrypDB-67 20240206, a “local” database that includes the sequences of recombinant and target proteins, antibodies used for pull-down experiments and common contaminants. Cysteine carbamidomethylation was set as a fixed modification and methionine oxidation and deamidation of glutamine and asparagine were considered as variable modifications; trypsin was specified as the proteolytic enzyme, with two missed cleavages allowed. The Mascot search results were imported into Scaffold (version 5.3.3, Proteome Software Inc., Portland, OR). A minimum of two identified peptides was required. The settings used resulted in a protein-level FDR of 0.3%. The top interactors of TcCARP3 were identified based on the enrichment of a protein in the TcCARP3-OE cell line eluate and absence in the EV eluate. Criteria for top interactors of TcCARP3 were infinite fold change (interacting protein present in all 3 of the TcCARP3-OE replicates and absent in the EV replicates analyzed), a total spectra count ≥ 1 in each of the 3 *TcCARP3*-OE replicates, and a P value < 0.05 after a t-test based on total spectra count, with Benjamini-Hochberg multiple test correction.

### Infection of kissing bugs with *T. cruzi* parasites

Kissing bugs (*Rhodnius prolixus*), were obtained from the colonies established at the Center for Disease Control (BEI Resources, NR-44077) (68). These kissing bugs were fed artificially bi-weekly with defibrinated rabbit blood (Hemostat) with a parafilm membrane feeding system (Hemotek). The colony condition was held at 24.0°C, 50 ± 10% relative humidity, and 6:00 am/6:00 pm light/dark photoperiods. Third instar *R. prolixus* were collected for infection with parasites. *T. cruzi* epimastigotes in exponential phase of growth were washed in 5 mL of 1x PBS pH 7.4 and mixed with defibrinated rabbit blood (complement inactivated at 56 ± 0.5°C before the addition of parasites) and offered to triatomines at 37°C through an artificial feeder at a concentration of 1×10^8^ parasites/mL (69-71). The kissing bugs were held at 24 ± 0.5°C, 50 ± 10% relative humidity, and 6:00 am/6:00 pm light/dark photoperiods to allow for parasite growth and differentiation. After four weeks, the hindguts were dissected out of the triatomine bugs, emulsified in 100μL of 1x PBS pH 7.4, and examined under the microscope for the presence of parasites in the hindgut to establish the percentage of infected insects. Three groups of 9-15 infected kissing bugs were dissected per *T. cruzi* cell line.

### Statistical analyses

Values are expressed as means ± Standard Deviation (SD). Statistically significant differences between treatments were compared using unpaired Student’s *t*-test, Kruskal-Wallis test, and one-way and two-way ANOVA tests with multiple comparisons, as mentioned in the legends of the figures. Differences were considered statistically significant for *P* < 0.05, and *n* refers to the number of independent experiments performed. All statistical analyses were performed using GraphPad Prism 9 (GraphPad Software, San Diego, CA).

## ACKNOWLEDGMENTS

Mass spectrometry analyses were conducted with expert technical assistance of Sammy Pardo and Dana Molleur at the University of Texas Health Science Center at San Antonio (UTHSCSA) Institutional Mass Spectrometry Laboratory, directed by Susan T. Weintraub, Ph.D. The laboratory is supported in part by UTHSCSA and by the University of Texas System Proteomics Core Network. We also thank Dr. Igor Almeida (University of Texas at El Paso, El Paso, TX) for his advice on the experimental design of mass spectrometry experiments. We thank Dr. Michael Boshart (Ludwig-Maximilians-Universität München, Germany) for antibodies anti-TbCARP3. We also thank Aqsa Raja for technical contribution in subcloning the *TcCARP3* gene in pTREXn-3xHA vector. Funding for this work was provided by the National Institute of Allergy and Infectious Diseases of the National Institutes of Health (Award number R00AI137322 to N. Lander and R21AI178573 to M.A. Chiurillo). Riley Hunter was an UPRISE program awardee at the University of Cincinnati. The funding agencies had no role in the study design, data collection and interpretation, or the decision to submit the work for publication. Opinions contained in this publication do not reflect the opinions of the funding agencies. We declare that we have no competing financial interests.

